# The Heterotaxy Gene *CCDC11* is Essential for Cytokinesis and Cell-Cell Adhesion via RhoA Regulation

**DOI:** 10.1101/452631

**Authors:** Saurabh Kulkarni, Rachel E. Stephenson, Sarah Amalraj, Ewelina Betleja, James J. Moresco, John R. Yates, Moe R. Mahjoub, Ann L. Miller, Mustafa K. Khokha

## Abstract

Mutations in CCDC11 have been identified in multiple patients with heterotaxy (Htx), a disorder of left-right (LR) patterning of the internal organs. In *Xenopus*, depletion of Ccdc11 causes defects in LR patterning, recapitulating the patient phenotype. Upon Ccdc11 depletion, normally monociliated cells of the Left-Right Organizer (LRO) exhibit multiple cilia per cell. Unexpectedly, we found that Ccdc11 is necessary for successful cytokinesis, and the multiciliation observed in Ccdc11-depleted cells was due to failed cytokinesis. Furthermore, CCDC11 depletion alters cell-cell adhesion with reduction in junctional localization of adhesion molecules. The small GTPase RhoA is critical for cytokinesis and cell-cell adhesion. Because the CCDC11 depletion phenotypes are reminiscent of RhoA loss of function, we investigated a possible connection to regulation of RhoA signaling. We demonstrate that CCDC11 is localized to the cytokinetic contractile ring overlapping with RhoA during cytokinesis and regulates total RhoA protein levels. Our results suggest that CCDC11 connects cytokinesis and LR patterning via RhoA regulation, providing a potential mechanism for heterotaxy disease pathogenesis.

## RESULTS AND DISCUSSION

### CCDC11 is important for establishing left-right asymmetry and proper cilia morphology in *Xenopus*

The embryo breaks bilateral symmetry beginning at the Left-Right Organizer (LRO) [1-3]. Here, a subset of cells with motile monocilia drive extracellular fluid towards the left, which is sensed by immotile monocilia[4-6]. As a consequence, *dand5* (Cerl2 in mouse, *coco* in frog), which is initially expressed bilaterally at the margin of the LRO, is repressed on the left [2, 7, 8]. Repression of *dand5* leads to expression of *pitx2* in the left lateral plate mesoderm [1, 9, 10]. Pitx2 drives asymmetric organogenesis such that the heart, which initiates as a midline tube, loops to the right while other organs like the gut also undergo asymmetric morphogenesis [11]. Therefore, a signaling cascade including cilia-driven extracellular fluid flow, *dand5*, and *pitx2* establish laterality of internal organs such as the heart.

CCDC11 plays a key role in establishing LR asymmetry [12-15]. A number of Htx patients have mutations in CCDC11, and depletion of Ccdc11 causes LR patterning defects in zebrafish and *Xenopus* [12-15]. However, analysis in *Xenopus* was restricted to examination of *coco* and *pitx2* [12]. To build on previous studies, we directly tested whether *ccdc11* is important for LR patterning in *Xenopus*. We depleted Ccdc11 using F0 CRISPR and examined cardiac looping (Figure 1A). Embryos depleted of Ccdc11 showed significant cardiac looping defects (~22%) compared to uninjected controls. In addition, overexpression of human CCDC11 also showed significant cardiac looping defects (~15%) compared to uninjected controls (Figure 1A), suggesting that the CCDC11 expression level is necessary for its proper function. To place Ccdc11 in the LR patterning cascade, we next examined *pitx2* and *coco*. Using a *ccdc11* morpholino oligo (MO), we found that *ccdc11* knockdown led to abnormal patterns of both *pitx2* (~25%) and *coco* (~80%) (Figures 1B-C). Therefore, using either depletion strategy, F0 CRISPR or MO, we show that *CCDC11* is critical for global LR patterning consistent with previous studies [12-14].

**Figure 1:**
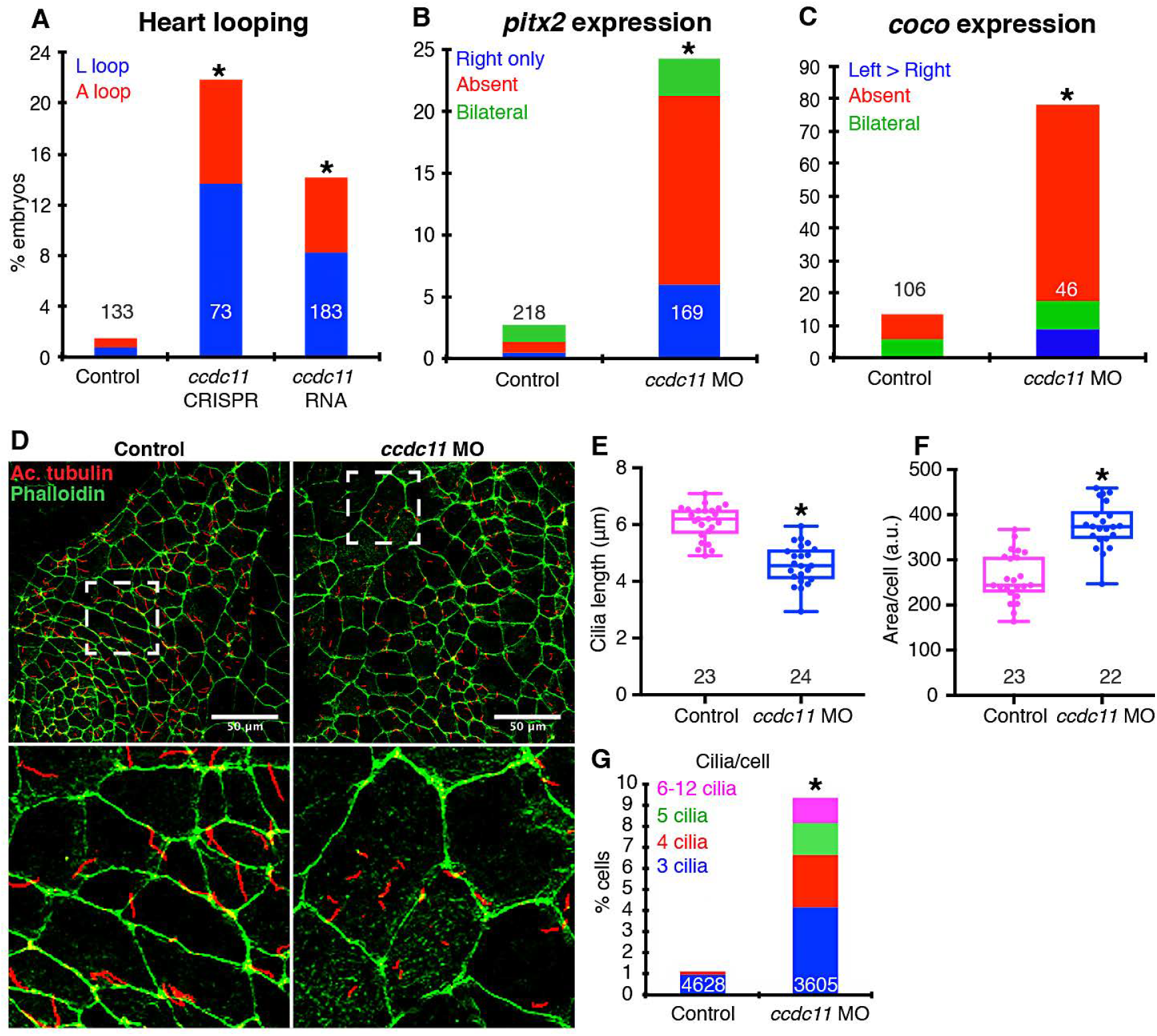
Ccdc11 depletion alters left-right patterning and cilia morphology in *Xenopus tropicalis*. (A, B, C) Percentage of *Xenopus tropicalis* embryos that have abnormal (A) looping of the heart outflow tube, (B) *pitx2*, or (C) *coco* expression. Embryos were injected at the one-cell stage with *ccdc11* CRISPR, *Xenopus ccdc11* mRNA, or *ccdc11* MO. L, outflow tube loops towards left; A, outflow tube fails to loop; Bilateral, signal present on both sides; Absent, signal absent on both sides; Reduced, signal reduced on both sides; Right only, signal is present on only right side; Left>Right, signal is more on the left side. ‘n’ = number of embryos. All results represent 3-4 independent experiments. (D) Cilia in control and *ccdc11* morphant (*ccdc11* MO) *Xenopus* LROs marked by anti-acetylated α-tubulin (red) and F-actin marked by phalloidin (green). Scale bar = 50 *μ*m. (E) Length of cilia in uninjected control and *ccdc11* morphant *Xenopus* LROs. Data is presented as box plot with 95% confidence interval. ‘n’ = number of embryos. Results represent 3 independent experiments. (F) Average cell apical surface area of uninjected control and *ccdc11* morphant *Xenopus* LROs. Data is presented as box plot with 95% confidence interval. ‘n’ = number of embryos. Results represent 3 independent experiments. (G) Percent of cells with multiple cilia in uninjected controls and *ccdc11* morphant *Xenopus* LROs. ‘n’ = number of cells. Results represent 3 independent experiments. ⋆= p < 0.001.

Previous studies have shown that CCDC11 is important for cilia, although the identified mechanisms differ [13, 14]. In the first study, CCDC11 appeared critical for cilia motility, specifically for docking of the outer dynein arm but not for ciliogenesis more generally [14]. On the other hand, a second study demonstrated that CCDC11 is critical for ciliogenesis [13]. Specifically, CCDC11 interacts with core components of the centriolar satellite proteins that are essential for ciliogenesis. This study also showed that loss of CCDC11 also causes defective assembly of cilia in multiciliated cells [13]. Based on these studies, we sought to examine cilia morphology in the LRO of *ccdc11* morphant *Xenopus tropicalis* embryos. The LRO is composed of monociliated cells where the cilium is normally ~5-6 *μ*m in length. Using immunofluorescence labeling, we found that cilia in *ccdc11* morphants were shorter (~4 *μ*m) (Figures 1D-E). Thus, consistent with the second study [13], our data suggest that Ccdc11 is important for general cilia structure. Interestingly, when we marked cell boundaries with phalloidin, which labels filamentous (F-) actin, we noted two additional striking findings. First, some of the Ccdc11-depleted LRO cells were dramatically larger compared to controls (Fig 1D,F). Second, many of these larger cells had multiple cilia per cell, in some cases as many as 10-12 cilia per cell (Figures 1D,G).

### CCDC11 is necessary for successful cytokinesis

We considered two hypotheses to explain why Ccdc11 depletion causes multiciliation in LRO cells: 1) Ccdc11 is important for regulation centriole biogenesis or 2) Ccdc11 is important for successful cytokinesis, and failed cytokinesis would lead to increased centriole numbers. When we labeled the nuclei of LRO cells, we found that cells with multiple cilia also had multiple nuclei (Figure S1), suggesting the Ccdc11 was important for cytokinesis. We quantified the multinucleation in Ccdc11-depleted *Xenopus tropicalis* embryos by labeling nuclei with H2B-RFP and marked the cell boundaries with phalloidin. When *ccdc11* was knocked down with a MO, approximately 30% of cells in stage 10 (early gastrulation) embryos had multiple nuclei (Figures 2A,B). When Ccdc11 was depleted using F0 CRISPR, approximately 20% of cells in the epidermis of stage 28 (post-neurulation) embryos exhibited multinucleation (Figures 2C,D).

**Figure 2:**
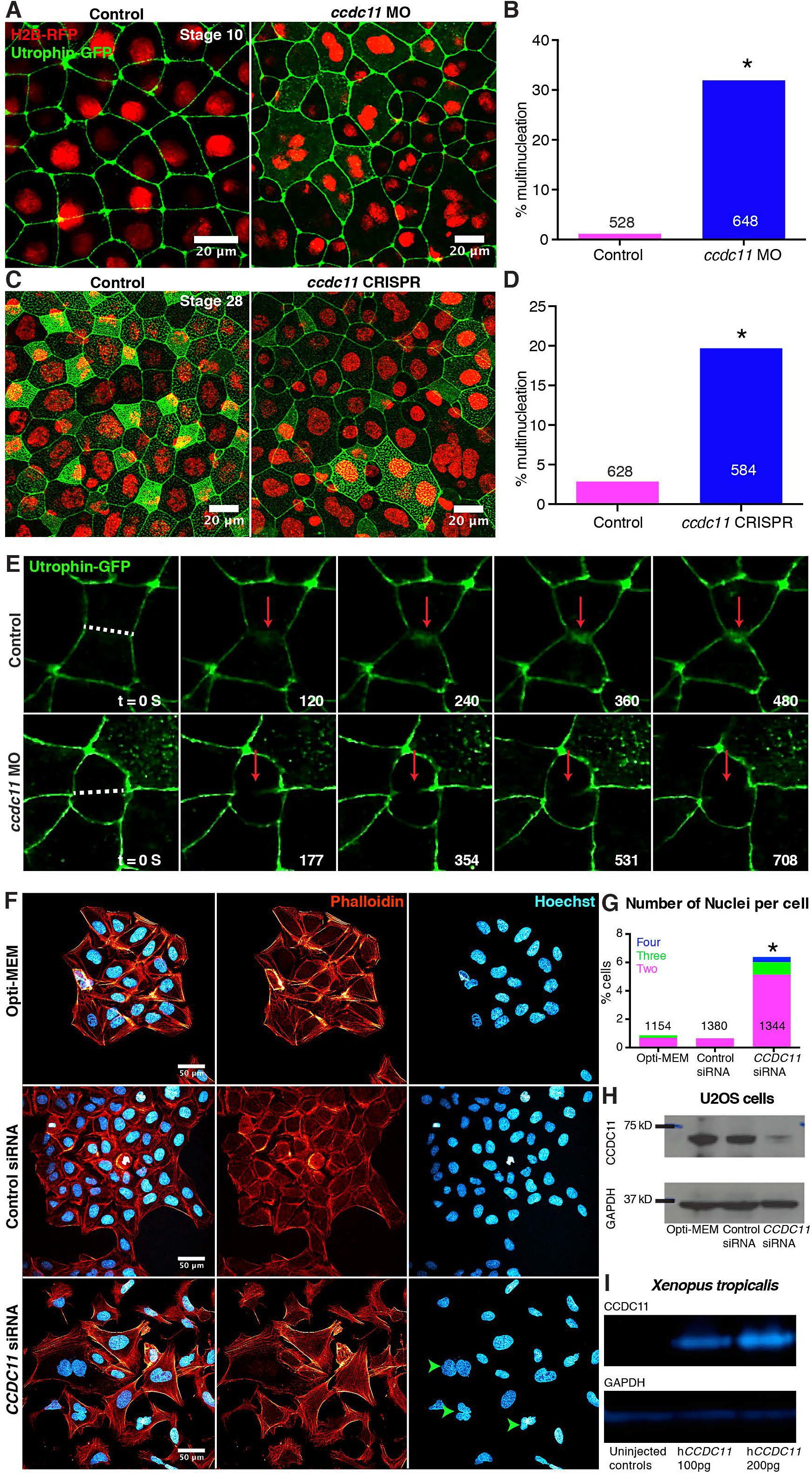
CCDC11 depletion leads to failed cytokinesis and multinucleation *Xenopus tropicalis* and human cells. (A-D) Percentage of cells with multiple nuclei in response to depletion of Ccdc11 by (A, B) *ccdc11* morpholino, or (C, D) *ccdc11* CRISPR. *Ccdc11* morpholino or CRISPR was injected in embryos at one cell stage. ‘n’ = number of cells. Scale bar = 20 *μ*m. Results represent 3 independent experiments. (E) Live imaging of *Xenopus tropicalis* embryo expressing utrophin calponin homology domain-(UtrCH-) GFP at Nieuwkoop and Faber stage 10 (pre-gastrulation) showing that cytokinesis is disrupted during actomyosin contractile ring ingression in *ccdc11* morphant. Results represent 2 independent experiments. (F, G) Percentage of human U2OS cells with multiple nuclei in response to CCDC11 depletion by siRNA. ‘n’ = number of cells. Scale bar = 50 *μ*m. Results represent 3 independent experiments. (H) Western blot showing loss of CCDC11 protein in human U2OS cells transfected with *CCDC11* siRNA. Results represent 2 independent experiments. (I) Western blot showing the antibody does not detect native *Xenopus tropicalis* Ccdc11 protein but can specifically detect human CCDC11 microinjected in *Xenopus tropicalis* embryos in a dose-dependent manner. Results represent 2 independent experiments. ⋆ = p < 0.001.

To visualize the spatiotemporal dynamics of cytokinesis, we performed live imaging in *Xenopus tropicalis* embryos. In control embryos, F-actin accumulated at the contractile ring during cytokinesis, and cytokinesis completed successfully. In contrast, cells depleted of Ccdc11 initiated cytokinesis, but failed to accumulate a significant amount of F-actin in the contractile ring, and failed to complete cytokinesis, even when monitored for longer durations (Figure 2E).

To test whether CCDC11 is also important for cytokinesis in human cells, we depleted CCDC11 using siRNA in human U2OS cells [13]. Indeed, *CCDC11* siRNA transfected cells showed significantly more multinucleation compared to cells transfected with control siRNA (Figures 2F-G). We confirmed CCDC11 depletion in U2OS cells using Western blot (Figure 2H). In order to test the specificity of the antibody, we overexpressed human *CCDC11* mRNA in *Xenopus tropicalis* embryos. The antibody does not detect native *Xenopus* Ccdc11 protein but does detect the human CCDC11 protein in a dose-dependent manner (Figure 2I).

### CCDC11 depletion affects cell-cell adhesion

While examining multinucleation in *CCDC11* siRNA-transfected U2OS cells, we noticed that cells did not appear to form proper cell-cell junctions; instead, the cells failed to pack and organize like controls. To examine if CCDC11 is important for cell-cell adhesion, we examined adherens junction molecules by immunofluorescence using antibodies against β-catenin and pan-cadherin (Figure S2A, C) [16]. In control cells, the β-catenin and pan-cadherin signal intensity peaked at the cell-cell junction with lower signal intensity in the cytoplasm (Figure S2A-D). However, in *CCDC11* siRNA transfected cells, both β-catenin and pan-cadherin signals were significantly reduced at cell-cell junctions (Figure S2A-D). Our results suggest that CCDC11 is not only important for successful cytokinesis but also for proper cell-cell adhesion.

### CCDC11 localizes to the contractile ring and midbody during cytokinesis

CCDC11, which is also known as cilia and flagella associated protein 53 (CFAP53), is known to localize to the ciliary base [13, 14]. However, given CCDC11’s newly identified role in cytokinesis and cell-cell adhesion, we decided to re-examine its localization. We began by examining ciliated cells including human retinal-pigmented epithelium (RPE) and pig kidney tubular epithelial cells (LLCPK). As expected, we detected CCDC11 at the base of cilia (Figures S3A-D), and consistent with a role in cytokinesis, CCDC11 was also localized to the midbody of recently divided cells (Figures S3A, C). To test antibody specificity, we depleted CCDC11 protein using siRNA in RPE cells and examined its fluorescence signal. siRNA depletion significantly reduced the CCDC11 signal both at the base of the cilium and the midbody demonstrating the specificity of the observed localization (Figures S4A-B).

While CCDC11 localizes to the midbody (Figures S3A, C), our live imaging of contractile ring ingression defects in Ccdc11-depleted *Xenopus tropicalis* embryos suggests that Ccdc11 may act at a much earlier stage in cytokinesis. Therefore, we tagged mNEON to the N-terminal of human *CCDC11*, injected the mRNA into *Xenopus laevis* embryos and examined CCDC11 localization using live imaging in the dividing cells of early (stage 5-7) embryos (Figure 3). We found that full length CCDC11 not only localized to the midbody but also localized to the contractile ring of dividing cells. In fact, CCDC11 localized to the contractile ring even before constriction initiated. These observations suggest that CCDC11 may act at the earliest steps of contractile ring ingression (Figures 2E and 3).

**Figure 3:**
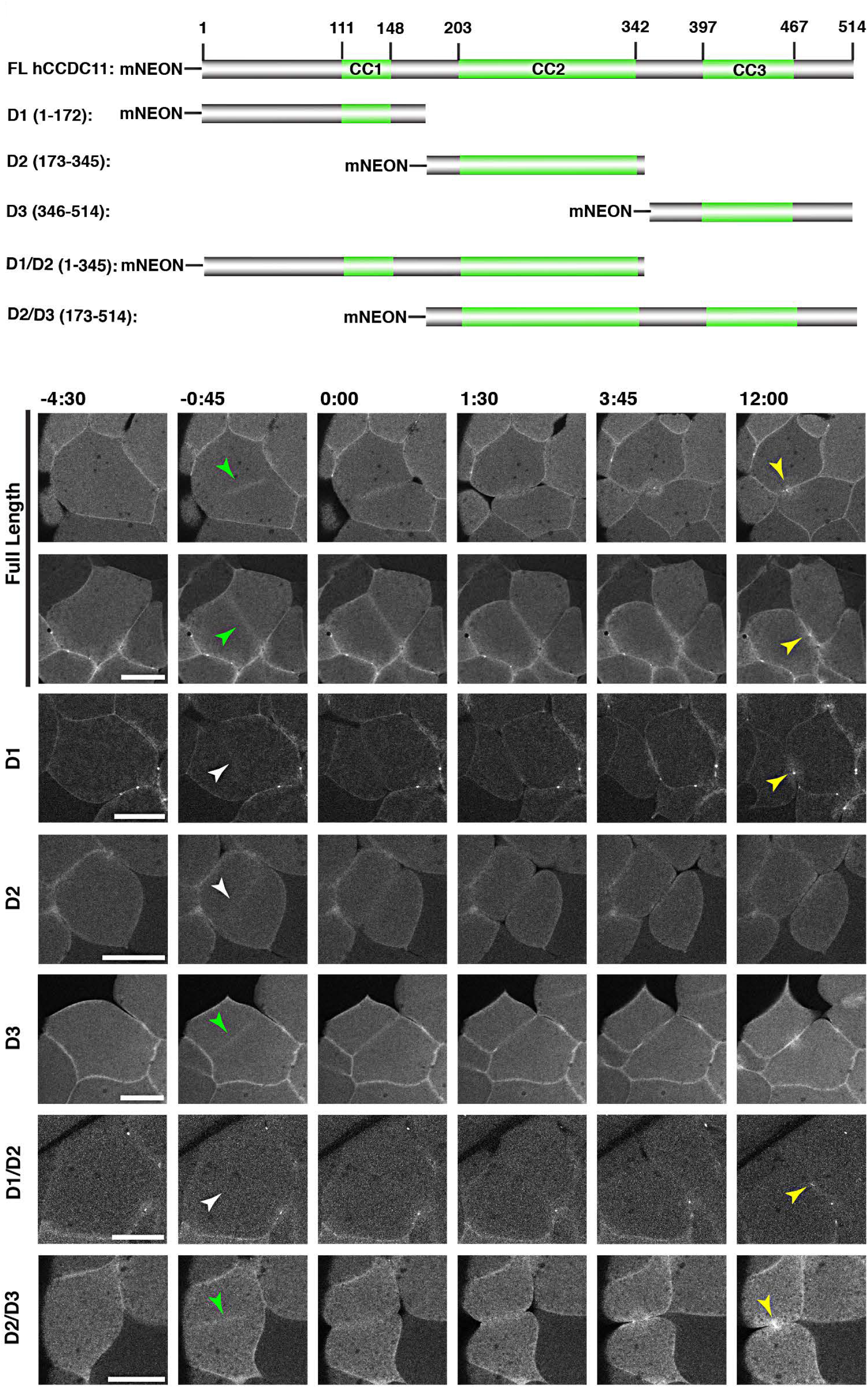
Domains important for Ccdc11 localization to the contractile ring and midbody during cytokinesis. Live imaging of mNEON-CCDC11 and fragments in blastomeres of *Xenopus laevis* embryos (Nieuwkoop and Faber stage 5-7). Time 0 is the time membrane ingression is first detectable. Arrowheads indicate the position of the contractile ring/midbody. Green arrowheads indicate strong accumulation of CCDC11 at the contractile ring, while white arrowheads indicate weak CCDC11 accumulation at the contractile ring. Yellow arrowheads indicate accumulation of CCDC11 at the midbody. Full length CCDC11 localizes to the cleavage furrow prior to ingression and the midbody following cytokinesis. D1 accumulates weakly at the contractile ring and strongly at the midbody. D2 accumulates weakly at the contractile ring. D3 accumulates strongly at the contractile ring. D1/D2 accumulates weakly at the contractile ring, indicating that D3 is necessary and sufficient for strong CR localization. D2/D3 accumulates strongly at the contractile ring and at the midbody, indicating that while D1 is sufficient for contractile ring localization, it is not required. Scale bar = 100 *μ*m.

### A C-terminal fragment of CCDC11 is necessary and sufficient for localization to the cytokinetic contractile ring

CCDC11 appears to be a multifunctional protein that localizes to and regulates different cytoskeletal structures including the ciliary base, cell-cell junctions, contractile ring, and midbody (Figures S2, S3, and 3) [13, 14]. CCDC11 contains three coiled coil (CC) domains [13]. However, whether any specific domains regulate differential localization is unknown. Because we were interested in understanding the role of CCDC11 in cytokinesis, we sought to identify the domain(s) critical for its localization to the contractile ring and midbody. To that end, we made five different fragments of human CCDC11 fused to mNEON: D1 (1-172aa), D2 (173-345 aa), D3 (346-514 aa), D1/D2 (1-345 aa) and D2/D3 (173-514 aa) (Figure 3). We injected the mRNA of each CCDC11 fragment and examined mNEON signal in dividing cells. D1, which contains the first CC domain, was weakly localized to the contractile ring towards the end of cytokinesis but was strongly localized to the midbody. D2 failed to localize to either the contractile ring or the midbody. D3 strongly localized to the contractile ring but weakly localized to the midbody. The D1/D2 construct localized similarly to D1, and the D2/D3 construct localized similarly to D3 suggesting that D2 is neither repressive nor instructive towards localization of CCDC11 to the contractile ring and midbody. Taken together our results demonstrate that while CCDC11’s first CC domain is necessary and sufficient for localization to the midbody, the third CC domain is necessary and sufficient for localization to the contractile ring during cytokinesis.

### CCDC11 co-localizes with RhoA during cytokinesis

To understand how CCDC11 regulates cytokinesis and cell-cell adhesion, we immunoprecipitated CCDC11 (using two different tags, GFP and Myc) and identified interacting proteins using liquid chromatography mass spectrometry (LC-MS/MS). Interestingly, a significant number of cytoskeleton proteins were highly enriched in our immunoprecipitated samples (Table S1). Gene ontology (GO) analysis identified significant enrichment for processes related to the actin cytoskeleton organization (6 out of top 10) (Table S2). Further, pathway maps identified processes related to Protein Kinase cAMP-dependent (PKA) and Rho GTPase-mediated cytoskeleton organization and integrin-mediated cell adhesion (Table S2).

In addition to defects in cilia, cells depleted of CCDC11 displayed two major phenotypes: cytokinesis failure and reduction in cell-cell adhesion proteins (Figures 2 and S2). Therefore, we sought to further investigate potential connections with Rho GTPase signaling, as Rho GTPases are master regulators of cytokinesis and cell-cell adhesion [17, 18]. Specifically, different Rho GTPases, such as RhoA, Rac1 and Cdc42, are critical for the initiation, maintenance and function of cell-cell adhesion [18-20]. Their activity is more compartmentalized during cytokinesis where Cdc42 is critical for mitosis, and RhoA is critical for cytokinesis [18]. Since we identified defects in cell-cell adhesion and cytokinesis when cells were depleted of CCDC11, we decided to investigate a role for CCDC11 in regulating RhoA. We began by examining localization of CCDC11 in relation to RhoA in dividing U2OS cells. We found that CCDC11 localization overlapped with RhoA localization both in the contractile ring and midbody supporting our hypothesis that CCDC11 may be affecting cytokinesis via manipulating RhoA activity (Figure 4A).

**Figure 4:**
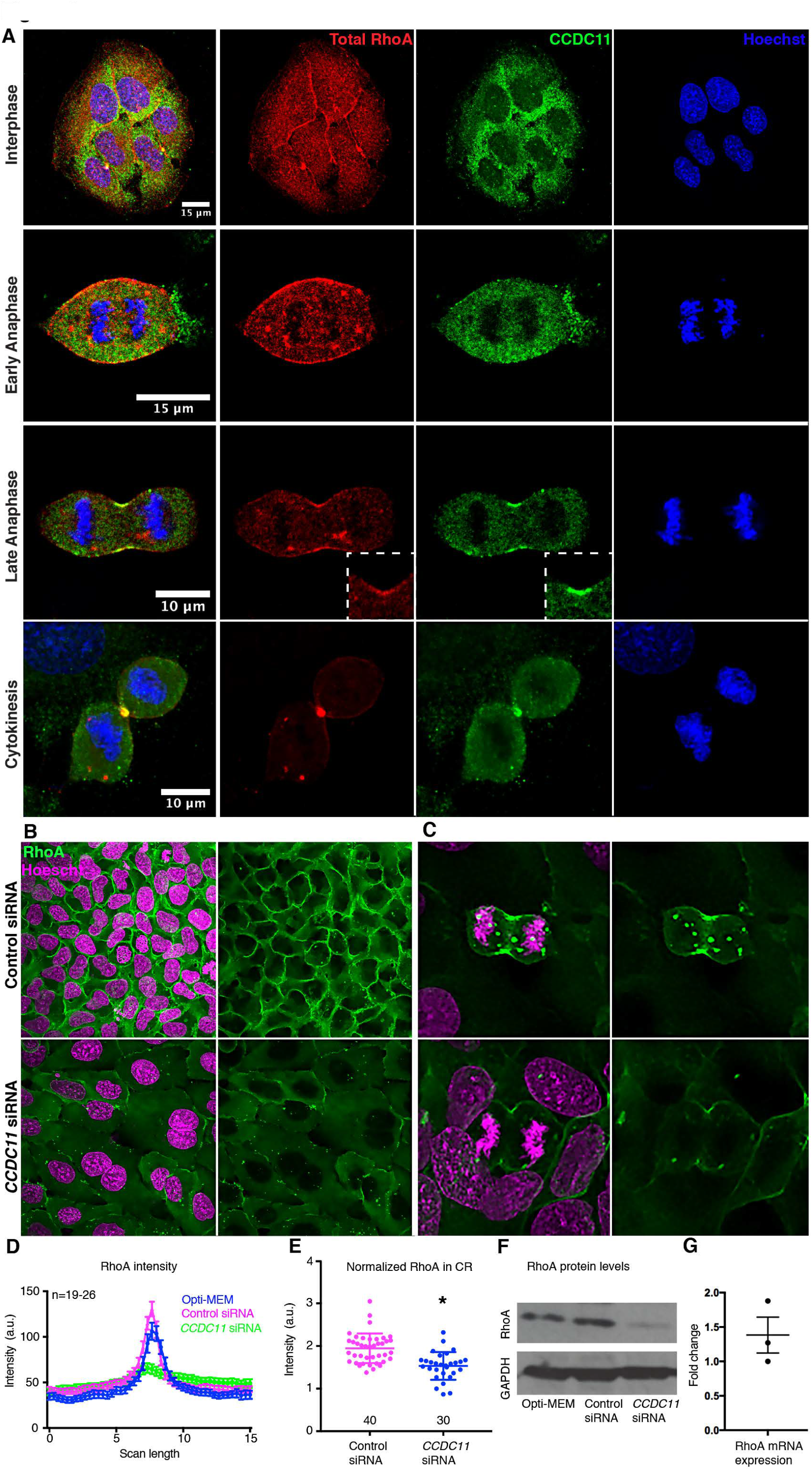
Ccdc 11 colocalizes with Rho A to the contractile ring and is important to maintain Rho A protein levels prost-transcriptionally. (A) CCDC11 co-localizes with RhoA in the contractile ring and midbody during cytokinesis in U2OS cells. Scale bars are 15 *μ*m for Interphase and Early Anaphase and 10 *μ*m for Late Anaphase and Cytokinesis. Results represent 2 independent experiments. (B, C, D, E) CCDC11 depletion affects RhoA accumulation at (B, D) cell-cell junctions, and (C, E) the contractile ring in the U2OS cells. The error bars represent standard error. ‘n’ = number of cells. Results represent 3 independent experiments. (F) Western blot showing reduction in RhoA protein in human U2OS cells transfected with *CCDC11* siRNA. Results represent 3 independent experiments. (G) Quantitative PCR data showing fold change mRNA levels of RhoA in *CCDC11* siRNA compared to controls. Results represent 3 independent experiments. ⋆ = p < 0.001.

### CCDC11 is essential for maintaining RhoA protein levels independent of transcription

To understand how CCDC11 might affect RhoA, we first examined RhoA localization at the cell membrane and contractile ring in CCDC11-depleted U2OS cells using TCA fixation, which better preserves the membrane-bound fraction of RhoA [21]. We found that CCDC11 depletion led to a significant reduction in RhoA signal at cell-cell junctions (Figures 4B, 4D). CCDC11 depletion also led to significant reduction of RhoA in the contractile ring (Figures 4C, 4E). To determine whether there was less RhoA protein or a relocalization of Rho away from the contractile ring in CCDC11-depleted cells, we performed Western blots for RhoA levels. In CCDC11-depleted cells, total RhoA protein levels were reduced (Figure 4F).

Loss of RhoA protein when *CCDC11* is knocked down suggested CCDC11 may regulate RhoA at the transcription level. To test the hypothesis, we compared RhoA transcript levels between control cells and CCDC11-depleted cells using quantitative PCR. In CCDC11-depleted cells, RhoA transcripts are in fact mildly elevated compared to controls demonstrating that CCDC11 does not regulate RhoA transcription (Figure 4G). This suggests that CCDC11 regulates RhoA post-transcriptionally, may be the stability of RhoA protein levels.

In summary, our results confirm previous findings that CCDC11 plays a key role in establishing organ laterality, a role conserved from fish to humans [12-15]. As evidenced by abnormal *coco* and *pitx2* expression, cilia-mediated signaling is aberrant in *Ccdc11*-depleted embryos. Previous studies suggested that CCDC11 is vital for ciliogenesis, and our results confirm defects in cilia length [13]. However, we discovered a “multiciliated” cell phenotype in the LRO that we traced to a defect in cytokinesis. We propose that depletion of Ccdc11 leads to cytokinesis defects in LRO cells, resulting in multiple nuclei and multiple centrioles. The presence of multiple centrioles gives rise to multiple cilia, which are structurally abnormal. Further, these cilia are not localized posteriorly in LRO cells indicating polarity defects that may contribute to abnormal left-right patterning. In addition, the defect in cilia signaling is amplified by the loss of CCDC11 from the base of the cilium, where it is required for composition of core components of satellite proteins [13]. Thus, the resulting LR patterning and cilia defects arise from a combination of CCDC11 function both in cytokinesis and in ciliary base structure.

A growing number of studies have shown a strong connection between cytokinesis and ciliogenesis, as both processes are actin and microtubule dependent [22-34]. Recently, a study showed that the midbody hosts multiple proteins important for cilia assembly. These proteins travel to the apical surface of a cell and move closer to the centrosome to enable cilia formation, providing a direct biological mechanism connecting these two important processes [23]. Interestingly, the first coiled coil domain of CCDC11 is required for localization to the centrioles, basal body and midbody [13]. Both results taken together provide some insights into how CCDC11 localization is regulated during the cell cycle. Meanwhile, others have identified large number of proteins that localize both to the contractile ring and the bases of cilia. In fact, the core cilia proteome and cytokinesis proteins overlap by more than 17 proteins [27]. For example, central spindle proteins essential for cytokinesis like PRC1, MKLP1 and INCENP are directly localized to the bases of cilia [27]. The master regulator of cytokinesis, RhoA, also localizes to the base of cilium [31-33]. Recently, we and others discovered that the chromatin modifier WDR5 is also localized to the bases of cilia, spindle microtubules, and the midbody [22, 24, 34]. Further, intraflagellar transport proteins necessary for ciliogenesis and cilia maintenance are also critical for cleavage furrow ingression [25, 26, 28-30]. Thus, localization and functional analysis of CCDC11 at the bases of cilia and at the cleavage furrow further highlights the importance of this group of multifunctional proteins that are essential in two critical biological processes: cytokinesis and ciliogenesis.

Cytokinesis failure can lead to tetraploidy or aneuploidy and potentially tumorigenesis [35, 36]. In fact defects in centrosome numbers are often a characteristic of cancer cells. For example, cancers of the breast, prostate, bladder, and pancreas are a subset that is often associated with centrosomal abnormalities [37]. Interestingly, CCDC11, which we show here is critical for cytokinesis, is implicated to play a role in cancer, specifically breast and prostate cancer [38, 39]. Further, in a few cases, an association between heterotaxy and cancer has been identified [40-42]; however, no clear mechanism emerged. Our study suggests that cytokinesis defects may represent a common underlying mechanism explaining the connection between heterotaxy and cancer. Finally, our results emphasize the importance of patient-driven gene discovery coupled with basic cell and developmental biology to discover new biological mechanisms. Identification of multiple roles for CCDC11 in cilia and cytokinesis may have important implications for patient management.

## ACKNOWLEDGMENTS

We want to thank Prof. Thomas Pollard on his valuable suggestions. We also want to thank Prof. Robert Denver on providing *Xenopus tropicalis* embryos. This project was funded in part by grants from the National Heart, Lung and Blood Institute K99/R00 (5K99HL133606) award to SSK, National Heart, Lung and Blood Institute (R01-HL128370) and National Institutes of Diabetes and Digestive and Kidney Diseases (R01-DK108005) to MRM, National Institute of Child Health and Human Development (R01HD081379) to MKK, National Institute of General Medical Sciences (P41 GM103533 to JRY), NIH grant R01GM112794 to ALM, and NSF Graduate Research Fellowship (DGE #1256260) to RES. MKK is a Mallinckrodt Scholar.

## METHODS AND MATERIALS

### Animal husbandry

*Xenopus tropicalis* were housed and cared for in our aquatics facility according to established protocols that were approved by the Yale Institutional Animal Care and Use Committee (IACUC). *Xenopus laevis* and *Xenopus tropicalis* were housed and cared for according to established protocols that were approved by the University of Michigan Institutional Animal Care and Use Committee (IACUC).

### Cell culture

Human retinal-pigmented epithelial (RPE) and pig kidney tubular epithelial cells (LLCPK) cells were obtained from Brueckner lab from Yale School of Medicine. U2OS cells were obtained from Carroll lab at Yale School of Medicine. Cells were cultured in DMEM/Ham’s F12 media supplemented with 10% fetal bovine serum (FBS).

### Microinjection of morpholino oligonucleotides and mRNA in *Xenopus*

*Xenopus tropicalis* embryos were produced by *in vitro* fertilization and raised to appropriate stages in 1/9xMR + gentamycin according to established protocols [43, 44]. Staging of *Xenopus* tadpoles was as previously described [45]. Morpholino oligonucleotides or mRNA were injected into one-cell or two-cell embryos as described previously [43]. The morpholino oligonucleotide for *ccdc11* translation blocking was injected at the concentration 7.5-10ng (5’-CATGCTTTCTCCCCAGCCGTGCTGT -3’). The CRISPR for *ccdc11* was injected at the concentration 800pg (5’-AAAGGAGCCGCCCCCGCCTT -3’). Alexa488 (Invitrogen) or green fluorescent protein (100pg) were injected as tracers when required. We generated *in vitro* capped mRNA using the mMessage mMachine kit (Ambion) and followed the manufacturer’s instructions. Full length *Xenopus ccdc11* image clone (IMAGE: 7730599) and human *CCDC11* image clone (IMAGE: 4831081) were subcloned into the pCS2+ vector using PCR amplification and cloning. Full length and mutant constructs for domain analysis were generated using PCR amplification and cloning into the pCS2+ vector with mNEON fluorescent protein. *Xenopus* WT CCDC11 was injected (100pg) at one-cell embryos for overexpression. *Xenopus laevis* embryos were injected with mRNA encoding mNEON-CCDC11 at following concentrations: full length (FL) 412.5 pg, the single domains (D1, D2, D3) 137.5 pg, and the doubles (D1/D2, D2/D3) 275 pg at the one cell stage. H2B-RFP and Utrophin-GFP were injected at 150pg at one cell stage of *X. tropicalis* embryos.

### siRNA in cell culture

For siRNA experiments, cells were transfected with *CCDC11* siRNA described before in [13] using Lipofectamine RNAiMAX transfection Reagent (Life Technologies) following the manufacturer’s instructions. U2OS cells with ~30% confluency were incubated with siRNA mix for 6 hours in the media without FBS. Cells were then transferred to the media supplemented with 10% FBS for 72 h before fixation. RPE cells with ~90% confluency were incubated with siRNA mix overnight in the media without FBS. The next day, cells were transferred to the new media without FBS for 72 hours to grow cilia.

### Cardiac looping in *Xenopus*

Frog embryos at stage 45 were treated with benzocaine and ventrally scored for cardiac looping using a light dissection microscope as previously described [46]. Loop direction is defined by the position of the outflow tract relative to the inflow of the heart: outflow to the right – D loop; outflow to the left – L loop; outflow midline, fails to loop – A loop.

### Image analysis

Images were captured using a Zeiss 710 Live confocal microscope except CCDC11 domain analysis imaging was done using Olympus Fluoview F1000 inverted confocal microscope. Images were processed in Fiji, Image J, or Adobe Photoshop. Quantification of GRP cilia length, cell number and LRO area was done using Velocity software on 3D image stacks. Number of cilia per cell in the *Xenopus* LRO was counted manually. Percent cells with multinucleation in *Xenopus* and U2OS cells were counted manually. Average apical surface area of LRO cells was calculated by measuring the total area of the LRO divided by the number of LRO cells, both marked for filamentous actin (phalloidin). Final figures were made in Adobe illustrator.

### Quantitative PCR

RNA expression levels were quantified using real-time quantitative PCR. Total RNA was isolated from 80-90% confluent U2OS cells treated either with control siRNA or *CCDC11* siRNA using RNaesy Plus Mini kKit (Qiagen). Three replicate RNA isolations were prepared independently for each treatment. Complementary DNA was generated from 3 *μ*g of total RNA by reverse transcription using iScript cDNA synthesis Kit (BIORAD) and IQ Supermix (BIORAD). Three technical replicates were performed. Gene specific primers were designed to amplify 100 – 150 bp fragments. To exclude genomic contamination, primers were designed spanning an intron. GAPDH was used for normalization of all data. Transcript abundance was quantified using the Applied Biosystems 7900HT. Differences in mRNA expression levels were determined using the standard curve method.

### Immunofluorescence

*Xenopus tropicalis* embryos at stage 16 were fixed in 4% paraformaldehyde/PBS overnight at 4°C to harvest LROs. Dissected LROs were then assayed for acetylated α-tubulin and phalloidin. U2OS cells were fixed in 4% paraformaldehyde/PBS for 2 hours at room temperature to assay, β-catenin, pan-cadherin with phalloidin and Hoechst. U2OS cells were fixed in 2% TCA (Trichloroacetic acid)/ PBS for 10 minutes and then washed with PBS to assay RhoA. RPE and LLCPK cells were fixed in 100% chilled methanol overnight at −20°C for γ-tubulin and CCDC11.

### Immunoprecipitation and Mass Spectrometry

For immnoprecipitation of GFP-or Myc-Ccdc11, RPE-1 cells stably expressing GFP-Ccdc11 or Myc-Ccdc11 at near endogenous levels[13] were grown on twenty 150-mm cell culture dishes to ~ 90% confluence, washed in PBS and trypsinized for 5 min in a 37°C incubator. After centrifugation, the cellular pellet (~ 1000-1500 *μ*L packed cell volume) was lysed in ice-cold Lysis Buffer (50 mM HEPES, pH 7.4, 1 mM MgCl_2_, 1 mM EGTA, 150mM NaCl and 10% glycerol) containing protease inhibitors (Mammalian Protease Arrest, G Biosciences), phosphatase inhibitors (PhosSTOP, Roche), and 1% NP-40 (Sigma) for 30 min on ice. Lysates were cleared by centrifugation at 3,000 g for 15 min, then at 50,000 × *g* for 30 min in an OPTIMA LE-80K ultracentrifuge using SW 41-Ti rotor (Beckman Coulter). Protein concentration of the clarified cellular extract was determined using a Pierce 660 nm Protein Assay kit. Equal amounts of the clarified cellular extract [~ 1 mg/mL] were mixed with 150 *μ*L of affinity resin composed of the following antibodies: 55 *μ*g goat anti-GFP (Rockland), 55 *μ*g control goat IgG (Santa Cruz Biotechnology), 20 *μ*g of rabbit anti-Myc (Sigma), or 20 *μ*g control rabbit IgG (Santa Cruz Biotechnology). All antibodies were covalently coupled to Protein-G or Protein-A Magnetic Beads via imidoester crosslinker (DMP; Pierce) per manufacturers protocol. Samples were incubated for 2 h at 4°C with rotation. The beads were magnetically pelleted and washed five times with ice-cold Lysis Buffer for 5 min each. Protein complexes were eluted in 240 *μ*L of Lysis Buffer.

MudPIT analysis of immunoprecipitated complexes were performed from protein pellets dissolved in buffer (8 M Urea 100 mM Tris pH 8.5) reduced with TCEP (Tris[2-Carboxyethyl]-Phosphine Hydrochloride), and alklyated with chloroacetamide. After dilution of urea to 2 M, proteins were digested with trypsin. Digested peptides were analyzed by LC/LC/MS/MS using an LTQ-Orbitap mass spectrometer. Multidimensional chromatography was performed online [47]. Tandem mass spectra were collected in a data-dependent manner with up to 10 ms2 scans performed for each initial scan (m/z range 300-2000). The search program Prolucid was used to match data to a human protein database [48]. Peptide identifications were filtered using the DTASelect program requiring two peptides per protein and a protein false discovery rate of 1% [49]. Interactors were selected as proteins that had more than five cumulative spectral counts in the experimental samples and were not detected in the controls.

### Statistical analysis

Sample size (n) is defined in every figure legend. Statistical analysis was done using PRISM and Vassarstats software. Heart looping, *pitx2c* and *dand5* were compared using Chi-square analysis. “Multiciliation” in the LRO cells, multinucleation in *Xenopus* and U2OS cells, and CCDC11 localization signal intensity in response to siRNA in RPE cells were also compared using Chi-square analysis. All other comparisons were made using t-tests after confirming the normal distribution of the data. We randomly picked one-cell *X. tropicalis* embryos from fertilization as uninjected controls or for morpholino oligo, CRISPR, or RNA injections.

### Antibodies

Mouse monoclonal Anti-acetylated α-tubulin (Sigma, T-6793; 1:1000) and Alexa 488 or Alexa 647-phalloidin (1:40) were used to mark the ciliary axoneme and F-actin respectively. Mouse monoclonal Anti-γ tubulin (Sigma, T6557; 1:100) was used for immunofluorescence. Rabbit polyclonal Anti-CCDC11 (Sigma, HPA041069) was used 1:100 for immunofluorescence and 1:500 for Western blot. Rabbit polyclonal Anti-CCDC11 (Sigma, HPA040595, 1:100) was used for immunofluorescence. Mouse monoclonal Anti-RhoA (Neweast biosciences, 26007) was used 1:50 was used for immunofluorescence and 1:1000 was used for Western blot. Rabbit polyclonal Anti-β-catenin (Santa Cruz, sc-7199, 1:100) and Rabbit polyclonal Anti pan-cadherin (Abcam, ab-6529, 1:100) was used for immunofluorescence.

